# The effects of sequencing depth on the assembly of coding and noncoding transcripts in the human genome

**DOI:** 10.1101/2022.01.30.478357

**Authors:** Isaac Adeyemi Babarinde, Andrew Paul Hutchins

## Abstract

**Background:** Investigating the functions and activities of genes requires proper annotation of the transcribed units. However, transcript assembly efforts have produced a surprisingly large variation in the number of transcripts, and especially so for noncoding transcripts. The heterogeneity of the assembled transcript sets might be partially explained by sequencing depth.

**Results:** Here, we used real and simulated short-read sequencing data as well as long-read data to systematically investigate the impact of sequencing depths on the accuracy of assembled transcripts. We assembled and analyzed transcripts from 671 human short-read data sets and four long-read data sets. At the first level, there is a positive correlation between the number of reads and the number of recovered transcripts. However, the effect of the sequencing depth varied based on cell or tissue type, the type of read considered and the nature and expression levels of the transcripts. The detection of coding transcripts saturated rapidly for both short-read and long-reads, however, there was no sign of saturation for noncoding transcripts at any sequencing depth. Increasing long-read sequencing depth specifically benefited transcripts containing transposable elements. Finally, we show how single-cell RNA-seq can be guided by transcripts assembled from bulk long-read samples, and demonstrate that noncoding transcripts are expressed at similar levels to coding transcripts but are expressed in fewer cells.

**Conclusions:** This study shows the impact of sequencing depth on transcript assembly. Sequencing read depth has a relatively minor impact on coding transcript assembly, but a major effect on the assembly of noncoding transcripts. This study highlights important factors to consider when deciding the sequencing read depths to be used for transcript assembly.

## Introduction

The genomic era has brought about a deep understanding of genomes [1]. The era which brought about the announcement of human [2], rat [3], mouse [4], chimpanzee [5] and multiple other genomes have led to an increased understanding of genome structures [6–8]. In addition, technological advancements have led to the genome-wide annotation and assessment of functional genomic elements. Of particular importance has been the identification of transcribed units, which have been described in multiple species [9–12], making it possible to investigate transcript functions.

The investigation of the functions of transcripts in a biological system requires accurate annotation of the transcripts. Transcript annotation is crucial because it reveals the structure of the transcripts and gives hints about their functions. Because genes have specific structures, several computational-based gene prediction tools have been published [13–15]. The strengths and weaknesses of these tools have been identified [16, 17]. However, prediction from a genome requires a comprehensive understanding of gene properties [18]. Therefore, transcript prediction tools often skip non-canonical genes and occasionally pick up erroneous signals. Additionally, technical advances in high-throughput sequencing have blurred the definition of a gene, as sequencing data has revealed that much of the genome is transcribed in one form or another. Indeed, these same technological innovations in sequencing have been applied to enhance transcript annotations [19–21]. Nonetheless, despite decades of research there remains considerable ambiguity on the full transcriptome [22–27].

Although various classes of transcripts are known, transcripts can be generally divided into two classes based on protein-coding ability [28–30]. While coding transcripts code for proteins, noncoding transcripts do not code for viable proteins [25,30,31], although there are suggestions that they can give rise to short-length peptides [32–34]. A class of noncoding transcripts at least 200 nucleotides long, termed long non-coding RNA (lncRNA), has attracted a lot of attention. Many tools can predict the coding ability of a transcript with high accuracy [35–39], and the properties unique to coding and noncoding transcripts have been described in several studies [29,35,36,40–42]. For example, coding transcripts tend to have higher expression levels in bulk samples, be longer, less likely to be localized to the nucleus, more stable, less tissue-specific, more evolutionarily conserved and contain fewer transposable element (TE)-derived sequences. Noncoding transcripts are the reverse for all of these properties. This suggests that coding and noncoding transcripts may have different levels of complexity for transcript assembly. In fact, unlike coding transcripts, the overlap between noncoding transcripts across databases is very low [18]. Indeed, the majority of the novel transcripts have been reported to be noncoding transcripts [26, 42].

The low expression, poor concordance between databases and abundance of TE-derived sequences in noncoding RNAs suggest that noncoding transcript assembly is more complicated. Therefore, it is likely that more reads are required to assemble noncoding transcripts, however the full influence of sequencing depth for short and long-reads has not been previously described. In this study, we retrieved 671 publicly available short-read bulk RNAs-seq datasets from a panel of cell types and tissues with various numbers of reads to investigate the impact of sequencing depth on the assembly of coding and noncoding transcripts. We also focused on 150 human pluripotent stem cell (hPSC) samples for which extensive bulk short-read and matching long-read and single cell-RNA-seq (sc-RNA-seq) data are available. We report that the relationship between the number of reads and the number of recovered transcripts varied based on cell or tissue type, the type of read considered and the nature and expression levels of the transcripts, with the noncoding transcripts consistently benefiting more from deeper sequencing. Overall, we find that sequencing read depth has a relatively minor impact on the detection of coding transcripts, but a major effect on the recovery of noncoding transcripts.

## Results

### Relationship between sequencing depth and transcript assembly across cell types and tissues

To understand the relationship between sequencing depth and transcript assembly, we assembled and studied the properties of transcripts from transcriptome data for 671 human samples with various sequencing depths (**Table S1**), ranging from the smallest (165,322 read pairs) to the largest (236,275,714 read pairs) and a mean and median read depth of 38,132,992 and 25,717,722 read pairs, respectively. The analyzed samples were all paired-end short-reads of which at least 70% aligned to the hg38 genome. We investigated the read depth at three levels. The first level is the total number of reads. The second level is the number of reads that mapped to the genome. The third level is the number of reads that were assigned to a transcript. The third level represents the number of reads that contribute to transcript assembly. We investigated the relationships between these three levels of read depth and numbers of transcripts assembled from the samples. The assembled transcript sets included both GENCODE-annotated and non-GENCODE transcripts. Our analyses showed that the number of transcripts retrieved is proportional to the sequencing depth, although the relationship is not linear (**Figure 1A**). Overall, Spearman’s correlation coefficient (rho) was 0.861. The transcript set from the assembly of each sample was then divided into coding and noncoding transcripts based on the score determined by FEELnc [35]. In almost all samples, more coding transcripts were found than noncoding transcripts. The correlation between the number of transcripts and sequencing depth for coding and noncoding transcripts were 0.822 and 0.840, respectively, reflecting a slightly higher correlation for noncoding transcripts. The analyses involving the number of mapped reads (**Figure S1A)** and the number of reads assigned to transcripts (**Figure S1B**) produced similar correlations. In fact, there is a high correlation between the total number of reads, number of mapped reads and number of reads assigned to transcripts (**Figures S1 C-E**).

**Figure 1.**
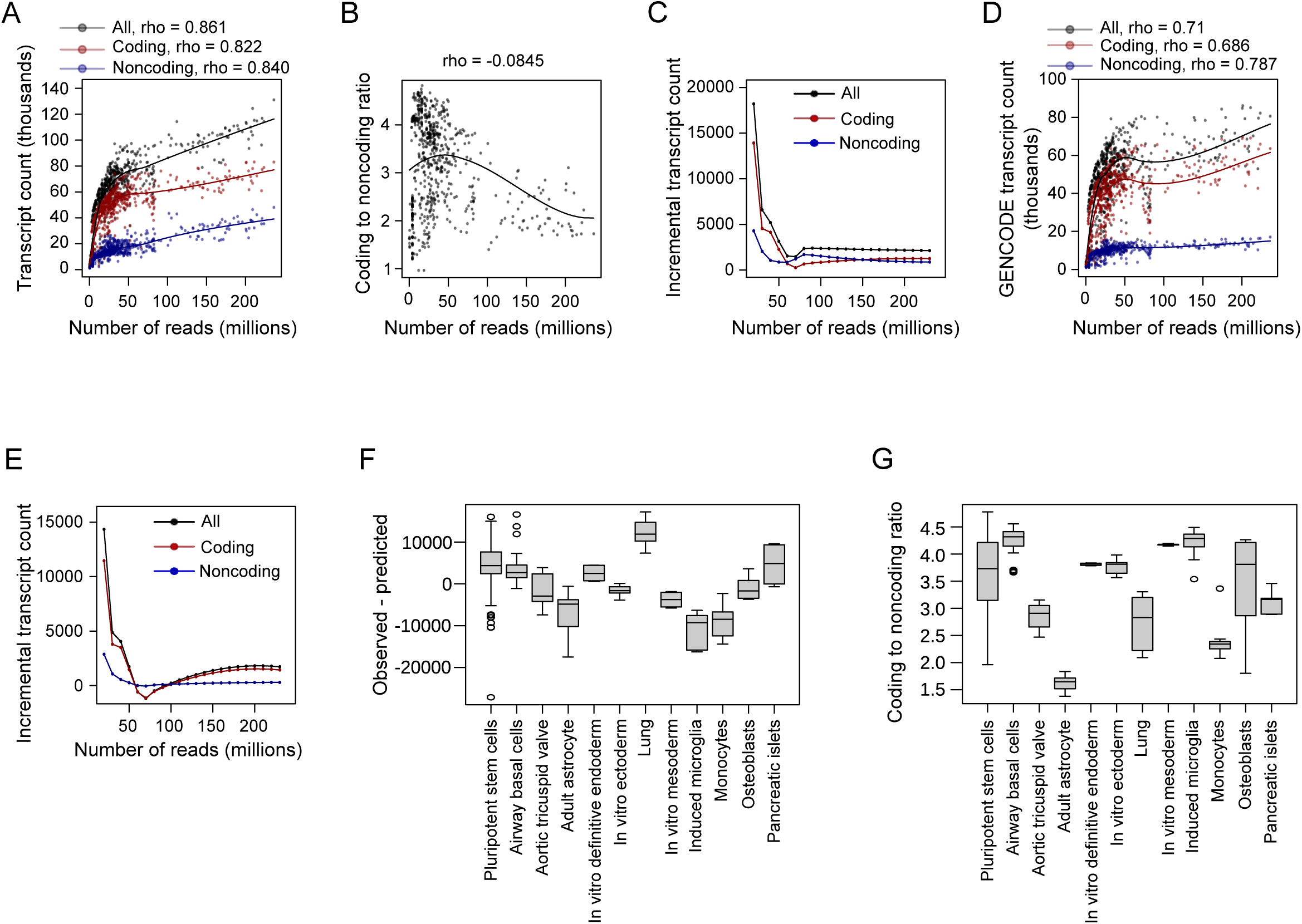
Relationship between the number of reads and assembled transcript count across multiple cells and tissues. Transcript assembly and coding potential evaluation were done for each of the 671 short-read samples from multiple cells and tissues. **A**. Number of reads versus the number of transcripts retrieved. For **panels A**, **B** and **D**, each point represents a sample while the line represents the loess fit. Rho is from Spearmans’ R. **B**. Number of reads versus coding to noncoding ratio. **C**. Increment in the number of transcripts per additional 10 million reads. **D**. Number of reads versus the number of transcripts for GENCODE-annotated transcripts. Assembled transcripts that were not found in GENCODE annotation were not included **E**. Increment in the number of transcripts per additional 10 million reads for GENCODE-annotated transcripts. **F**. The difference between the observed and the predicted numbers of transcripts based on the loess prediction of all the 671 samples. Cells or tissues with at least 8 biological or technical replicates in the dataset are presented. **G**. The coding to noncoding ratio varied across cells and tissue types. Only cells or tissues with at least 8 biological or technical replicates are shown.

The slightly higher correlations between noncoding transcripts and the sequencing depth suggest that the proportion of coding transcripts might be different across sequencing depths. Indeed, samples with relatively shallower depths tended to have higher coding to noncoding ratios (**Figure 1B**). This indicates that coding sequence transcripts saturate at relatively shallow sequencing depths. The polynomial fit shows that the ratio slightly increases initially, and starts dropping at around 50 million reads. For example, the ratio was around 5 for multiple shallow-depth samples. For all the samples with at least 200 million reads, the ratio was lower than 3, which suggests that deeper sequencing benefits noncoding transcripts.

We next checked for the increase in the number of transcripts for every 10 million additional reads. For these analyses, the predicted polynomial lines of best fit were used to predict the number of transcripts for each increase in reads. The result shows that the incremental number of transcripts was higher at shallow depth than at high depth (**Figure 1C**). The coding transcripts reached the lowest level of increase at 70 million reads. Essentially, this means that coding transcripts are accurately assembled from a relatively small number of reads. After 100 million reads, the incremental numbers of the transcripts remained relatively similar. These results may reflect differences in the number of noisy transcripts assembled that are specific to each sample. We therefore focused the analyses on GENCODE annotated transcripts only. Interestingly, the results remain essentially the same, with noncoding transcripts having a higher correlation between the number of retrieved GENCODE transcripts and the number of reads (**Figure 1D**). However, the correlations were generally lower (0.686 for coding, 0.787 for noncoding). The assembly of transcripts with increasing numbers of reads was also similar (**Figure 1E**), with more additional transcripts assembled at a relatively shallow sequencing depth. These results show that the relationship between the number of reads and the number of assembled transcripts is not just due to the inclusion of more transcriptional noise, but reflects the ability to retrieve more (potentially) genuine transcripts.

The inclusion of samples from different cell and tissue types in building the model inherently assumes that transcript abundance and coding proportion are similar across cell types. However, it is known that the number of transcripts can vary across tissues [43]. To explore this, we selected the 12 cells or tissues that had at least eight biological replicates from our dataset. We checked the difference between the actual number of transcripts assembled and the number of transcripts predicted from the read depth using the model in **Figure 1A**. Our analyses show that the difference between the observed and predicted number of transcripts varied across cells and tissue types (**Figure 1F**). Whereas some tissues have positive observed-predicted differences, indicating that the number of transcripts retrieved was more than the number predicted, others have negative observed-predicted differences, indicating that the number of transcripts retrieved was lower than the predicted value. Interestingly, this heterogeneity was also observed for both coding (**Figure S1F**) and noncoding transcripts (**Figure S1G**). Intriguingly, there was a rough inverse relationship between the observed-predicted transcript counts between coding and noncoding transcripts (**Figure S1F, G**), indicating that the ratio between coding and noncoding transcripts is tissue-specific (**Figure 1G**). Indeed, there was variation in coding to noncoding ratios across tissues, with the ratio ranging from ∼1.5 in adult astrocytes to ∼4.4 in airway basal cells (**Figure 1G**). These results suggest that the relationship between read depth and the number of retrieved transcripts varies across cells or tissues, and that including multiple tissues or cells may confound the results. The results also indicate that the coding to noncoding ration is not consistent, but varies substantially across different cell types and tissues.

### Impacts of sequencing depth on human pluripotent stem cell transcript assembly

We then decided to analyze a single cell type in depth. For this, we focused on human pluripotent stem cells (hPSCs). There are widespread problems in metadata annotations of sequencing samples [44–46], hence we used a set of 150 RNA-seq samples that we previously computationally verified to be normal undifferentiated hPSCs [42]. We also used the transcriptome assembly from that study, which was built using a combination of long and short-read RNA-seq data. The assembly from that study comes in two forms, an unfiltered set, containing lower confidence transcripts for a total of 272,209 transcripts, and a more confident filtered transcript set containing 101,492 transcripts, including 7,261 novel and 19,575 variant transcripts. Novel transcripts were defined as transcripts that do not overlap any known GENCODE transcript while variant transcripts were different isoforms of a GENCODE gene [42].

We investigated the effect of read depth on the number of transcripts retrieved in hPSCs. While the relationship between transcript count and the number of reads was generally higher than in the multi-sample analyses (**Figure 2A)**, a higher noncoding correlation (0.892 compared to coding’s 0.806) was still found. Similar results were also found in the relationship between transcript counts and the number of mapped reads (**Figure S2A**) or the number of reads assigned to transcripts (**Figure S2B)**. Unsurprisingly, there is a high correlation between the number of reads, the number of mapped reads and the number of reads assigned to transcripts (**Figures S2C-E**). Interestingly, the relationship between the coding to noncoding ratio and the number of reads was more obvious, and deeper sequenced samples tended to have a higher proportion of noncoding transcripts (**Figure 2B**). In fact, the coding to noncoding ratio of the deepest sample was around 2, compared to 4.7 in some shallow-depth samples. The incremental numbers of transcripts per increasing reads was also different between coding and noncoding (**Figure 2C**). At around 150 million reads, the incremental number of coding transcripts found was zero, suggesting that sequencing had reached full saturation for coding transcripts. However, for noncoding transcripts, even at 230 million reads (the largest single library in our data set), the incremental number of assembled transcripts did not reach zero, indicating that noncoding transcripts continued to benefit from deeper sequencing.

**Figure 2.**
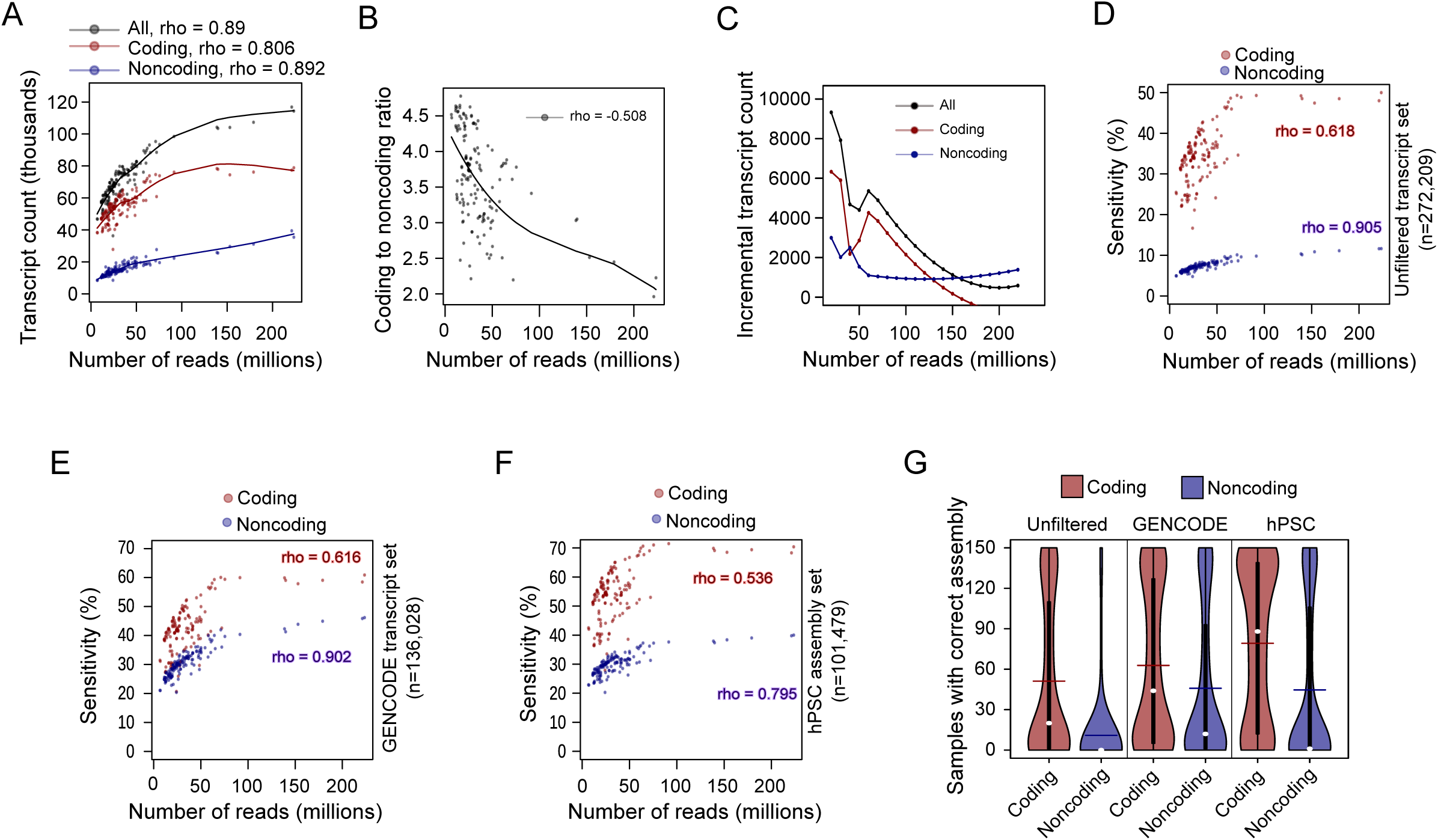
Sequencing depth is positively correlated with transcript count and sensitivity in human pluripotent stem cells. **A**. Number of reads versus the number of transcripts assembled from 150 computationally verified hPSC samples. The line indicates the loess fit, for this and in panel B. Rho is from Spearmans’ R. **B**. Number of reads versus the coding to noncoding ratio. **C**. Increment in the number of transcripts per additional 10 million reads. **D**. Sensitivity versus the number of reads using unfiltered transcript set assembled from the merged alignment of all the 150 hPSCs. **E**. Sensitivity versus the number of reads using only GENCODE-annotated reference. The hPSC transcripts that were not found in GENCODE annotation were excluded. **F**. Sensitivity versus the number of reads using hPSC assembly. The hPSC assembly, published in Babarinde *et al.* includes additional transcripts that were not found in GENCODE annotation but excludes GENCODE transcripts that are not expressed in hPSCs. **G**. The number of samples in which the transcripts are assembled. For each category of transcripts, the number of samples for which the transcripts were correctly assembled was computed.

We then set out to explore the sensitivity of the transcript assembly. We defined false positives as a transcript that matched to the unfiltered superset of hPSC transcripts (n=272,209), which were obtained by combining all 150 hPSC samples [42]. We checked the sensitivity of each sample in terms of the percentage of the transcripts that were correctly assembled, relative to the hPSC unfiltered superset. The sensitivity, computed per sample, reflects the percentage of the transcripts in the unfiltered set that were assembled in the sample assembly. Our result showed that the sensitivity of coding transcripts is consistently higher than that of noncoding transcripts (**Figure 2D)**. The sensitivity of coding transcripts seemed to reach a peak of about 50%, after 70 million reads. For all the samples, the sensitivity of noncoding transcripts never exceeded 15%. As expected, deeper samples have higher sensitivity. These differences between coding and noncoding transcripts prompted us to investigate the distribution of coding and noncoding transcripts in filtered and unfiltered transcript sets, as well as in the GENCODE transcript set. The unfiltered transcript set contained a higher proportion of noncoding transcripts than filtered transcript (**Figure S2F)**. This suggests that the filtering process disproportionately affected noncoding transcripts.

Since the unfiltered transcript set likely contains many unreliable transcripts, we repeated the hPSC transcript assembly sensitivity analysis by focusing only on a subset of unfiltered hPSC transcripts that are annotated in GENCODE (**Figure 2E)**. The patterns were essentially the same. Noncoding transcripts tended to have lower sensitivity but higher correlations between sensitivity and the number of reads. However, using the GENCODE transcript set as a reference increased the sensitivity, with the sensitivity of coding transcripts of some samples going above 60% and the noncoding sensitivity of some samples reaching 43%. Previous studies have reported that some genuine transcripts are missing in the GENCODE annotation [25,42,47,48]. Therefore, we repeated the analyses using the hPSC filtered transcript set [42]. The results remain essentially the same, except that the sensitivity of coding transcripts goes up, and the correlations became lower (**Figure 2F**). Because one of the steps in the hPSC transcript filtering process is the number of samples in which a transcript is expressed [42], we checked the number of expressing samples and found that indeed coding transcripts tended to be more broadly expressed (**Figure 2G**). The filtered coding transcript set tends to be more broadly expressed than even the full GENCODE transcript set (**Figure 2G**). These results show the relationships between the number of reads, the number of transcripts retrieved and the sensitivity, across various reference transcript sets. Whilst coding transcripts saturate rapidly at relatively low numbers of reads, the number of noncoding transcripts continues to increase with sequencing depth. However, the sensitivity of coding transcripts tends to be higher than that of noncoding transcripts, suggesting the extra noncoding transcripts might be less trustworthy.

### Effects of read depth and replication numbers on transcript assembly

The results in **Figures 2A-G** involved hPSC samples from different sources. The results might therefore reflect the effects of the number of reads or effects on the samples from multiple factors independent of sequencing depth. To investigate the impact of the number of reads on transcript assembly from homogenous background, we took a theoretical approach, and simulated 20 replicates each of 10 million paired-end reads, 10 replicates of 20 million reads, four replicates of 50 million reads, two replicates of 100 million reads and one replicate of 200 million reads. The simulation was done using *rsem-simulate-read* in the RSEM package [49]. Two samples, SRR597912 and SRR597895 [50], were used to build the reference upon which the simulation was generated, as these two samples together had about 460 million paired end reads. We first assembled transcripts for the merged reads of the two samples. We then built the model of the RNA-seq data using the assembled transcript set and the short-read RNA-seq data. Finally, we simulated different numbers of reads using the read simulator in RSEM [49]. Individual transcript assembly confirmed that the number of transcripts increases with the number of reads, and the fraction of coding transcripts tended to dominate in the total number of transcripts (**Figure 3A**). Similarly, the coding to noncoding ratio tended to decrease with sequencing depth confirming that deeper sequenced samples can indeed retrieve more noncoding transcripts (**Figure 3B**). The sensitivity of the assembly was also confirmed to increase with the sequencing depth, with the sensitivity in coding transcripts consistently higher than the sensitivity in noncoding transcripts (**Figure 3C**). Interestingly, the precision of the coding and noncoding transcripts for lower depths were similar (**Figure 3D**), suggesting that the increase in sensitivity is not always accompanied by a decrease in precision.

**Figure 3.**
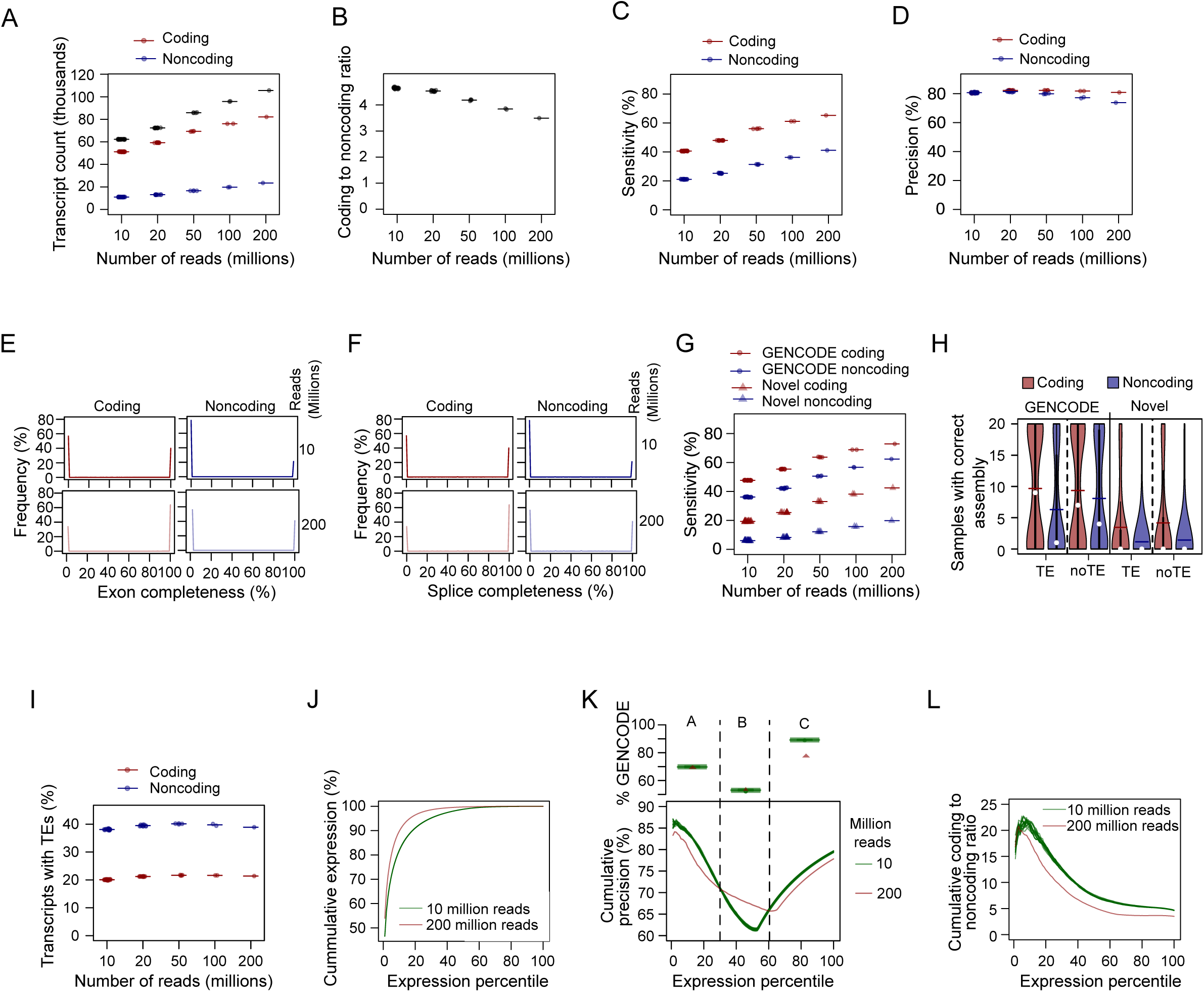
Effect of sequencing depth on the numbers and correctness of transcripts assembled from simulated short-read data. 10 million (n=20), 20 million (n=10), 50 million (n=4), 100 million (n=2) and 200 million (n=1) hPSC reads were simulated using SRR597912 and SRR597895 reads. Transcript assembly and coding potential evaluation were done for each simulated read sample. The relationship between the number of reads and transcript counts **(panel A)**, coding to noncoding ratio **(panel B)**, sensitivity **(panel C)** and precision **(panel D)** are presented for each sample. The transcript assembly obtained from the merged SRR597912 and SRR597895 reads was used as the reference for the estimation of sensitivity, precision and completeness. Exon (**panel E**) and splice (**panel F**) completeness of the reference coding and noncoding transcripts in different assemblies. The distributions are shown for coding and noncoding transcripts in 200 and 10 million read assemblies. **G**. Sensitivity is positively correlated with the number of reads. Samples with the same read numbers produced similar numbers of transcripts. **H**. The numbers of samples for which different categories of transcripts were correctly assembled. The assemblies of 20 replicates of 10 million reads were investigated to highlight correctly assembled transcripts. **I**. The proportion of TE-containing transcripts was similar across various read depths. **J**. Cumulative expression of assembled transcripts. The transcripts were ranked based on expression level, and the cumulative expression is presented in percentages. **K**. Cumulative precision of transcripts based on the expression levels. Based on the expression levels and the cumulative precision in 200 and 10 million read assemblies (lower panel), the transcripts were grouped into three classes. The upper panel shows the percentages of GNCODE-annotated transcripts in the three classes of transcripts. **L**. Cumulative coding to noncoding ratio varied with expression levels and the numbers of reads.

Next, we asked whether multiple replicates or single replicates of the same sequence depth would affect the quality of assembly. For this, we merged the alignments of 20 replicates of 10 million reads to make a total of 200 million reads. A similar procedure was carried out for 20, 50 and 100 million read simulated data such that all the categories summed up to 200 million reads. The results showed that there were no substantial differences in sensitivity (**Figure S3A**) and precision (**Figure S3B**) of the assembly. This result from stimulated sub-sampled data from a homogenous superset of reads suggests that the overall read depth is more important for assembling transcripts than the number of replicates.

The ‘correctness’ of a transcript assembly can also be assessed in terms of transcript completeness at the exon or splice level [42]. A reference assembly with 100% exon completeness has all its exons retrieved in the new assembly, whilst an incomplete assembly might have matching exons, but none of the exon/intron boundaries would match. Transcript completeness was compared between assemblies from 10 million reads (low depth) and assembly from 200 million reads (high depth). **Figure 3E** shows that the exons of each assembled transcript were mostly either found (100% exon completeness) or were completely missing (0% exon completeness), reflecting the integrity of the assembly. Partial exon overlaps were (surprisingly) not common. Consistent with the results in **Figure 3C**, the 200 million deep read assembly was more complete, and coding transcripts had higher exon completeness than noncoding transcripts. A valid transcript should have all the introns spliced out. So, it is important to investigate if all the splice sites are included. We computed splice completeness and found a pattern similar to exon completeness, with coding transcripts and the assembly from 200 million reads having higher overall splice completeness (**Figure 3F**). These results highlight the importance of read depth and the difficulty in assembling noncoding transcripts.

The reference transcripts used for the simulation comprised the GENCODE annotated transcripts as well as new transcripts that are not annotated in the GENCODE assembly, which we referred to as novel or variant in Babarinde *et al*. (2021). As previously reported [42], novel transcripts contained a higher percentage of noncoding transcripts (**Figure S3C**). Across both the GENCODE and novel transcript sets, coding transcripts tended to have higher expression levels (**Figure S3D**). Interestingly, the expression levels of novel transcripts tended to be higher than that of GENCODE annotated transcripts in both coding and noncoding transcripts. The results further highlight the fact that noncoding transcripts are more difficult to assemble and are depleted in GENCODE annotations.

We next checked if the sensitivity of the transcript assembly is similar between novel and GENCODE transcripts. Across different read depths, the GENCODE annotated set consistently had higher sensitivity (**Figure 3G**), suggesting that the novel transcripts are more difficult to assemble. One reason may be because of the higher proportion of transposable element (TE) fragment sequences in novel transcripts. Indeed, the presence of a TE in coding and noncoding transcripts reflects the assembly sensitivity in GENCODE and novel transcript sets (**Figure S3E**). Specifically, the GENCODE coding transcript set with the highest assembly sensitivity had the lowest TE content while the novel noncoding transcript set with the lowest assembly sensitivity contained the highest proportion of TE-containing transcripts. We next checked how the number of expressing samples is affected by TE presence and coding ability in GENCODE and novel transcript sets (**Figure 3H**). This analysis was done using 20 replicates of 10 million reads. GENCODE annotated transcripts tended to be more broadly expressed than novel transcripts, and coding transcripts tended to have broader expression than noncoding transcripts (Figure 3H). Also, transcripts with no TEs tended to be more broadly expressed, especially in GENCODE annotated noncoding transcripts. Interestingly, increases in sequencing depth do not seem to substantially improve the proportion of TE-containing transcripts in the assembly (**Figure 3I**), suggesting a complex impact of TE presence in transcript assembly from short-read data.

Next, we investigated the impact of sequencing depth on the expression levels of detected RNAs. First, we investigated the cumulative RNA levels in shallow-depth samples (10 million reads) and a high-depth sample (200 million reads). The assembly from 200 million reads reached saturation more quickly than the assemblies of 10 million reads (**Figure 3J**). For example, 95% of all detected RNAs from the high depth sample were found in the top 17% of highly expressed transcripts. On the contrary, the top 30% of the expressed transcripts in the shallow samples contributed 95% of all expressed RNAs. The investigation of transcript precision showed that transcripts can be classified into three groups based on the expression-ranked assembly precisions of shallow-depth (10 million read) and high-depth (200 million read) samples (**Figure 3K)**. Groups A (top 30 percentile in expression) and C (lowest 40 percentile in expression) transcripts have higher precisions in the shallow-depth samples while group B (30-60 percentile in expression) transcripts have higher precision in the high-depth sample. Group B transcripts have the lowest percentage of GENCODE transcripts suggesting that a substantial proportion of the novel transcripts have intermediate expression levels. Interestingly, there are no major differences in the percentages of GENCODE transcripts in groups A and B across low-depth and high-depth samples. However, group C transcripts have a higher GENCODE percentage in low-depth than high-depth samples. Coding to noncoding ratio falls very rapidly with expression (**Figure 3L**), highlighting that the expression level considered can affect the ratio observed. For example, if the top 10% of transcripts were considered, the set would have a similarly high coding to noncoding ratio in 200 million reads as in the 10 million read assemblies. The coding to noncoding ratio is reduced if the top 20% of the transcripts are considered, and the ratio would be lower in 200 million than 10 million read assemblies (**Figure 3L**). These results demonstrate that the effect of read depth on the assembly varies with the transcript expression level. Consequently, the composition of the transcripts is drastically altered by setting the expression threshold.

### Effect of sequencing depths on transcript assembly in long-read data

Recent studies have demonstrated the advantages of long-read sequences over short-read sequences for genome and transcript assembly [21,41,51,52]. We therefore investigated the effect of sequencing depths on long-read sequences from the PacBio platform. We retrieve the long-read data from H9 (∼25 million reads) cells and iPSCs (∼7 million reads). Transcript assembly from H9 returned 94,205 transcripts, including 74,163 coding transcripts while 59,254 transcripts, including 47,902 coding transcripts were assembled from iPSCs (**Figure 4A**). These corresponded to a 3.7 and 4.2 coding to noncoding ratio in H9 and iPSCs, respectively. Although only two datasets were considered, the observation of a lower coding to noncoding ratio in the deeper read assembly was confirmed, as observed in short-read data (**Figures 2B, 3B**).

**Figure 4.**
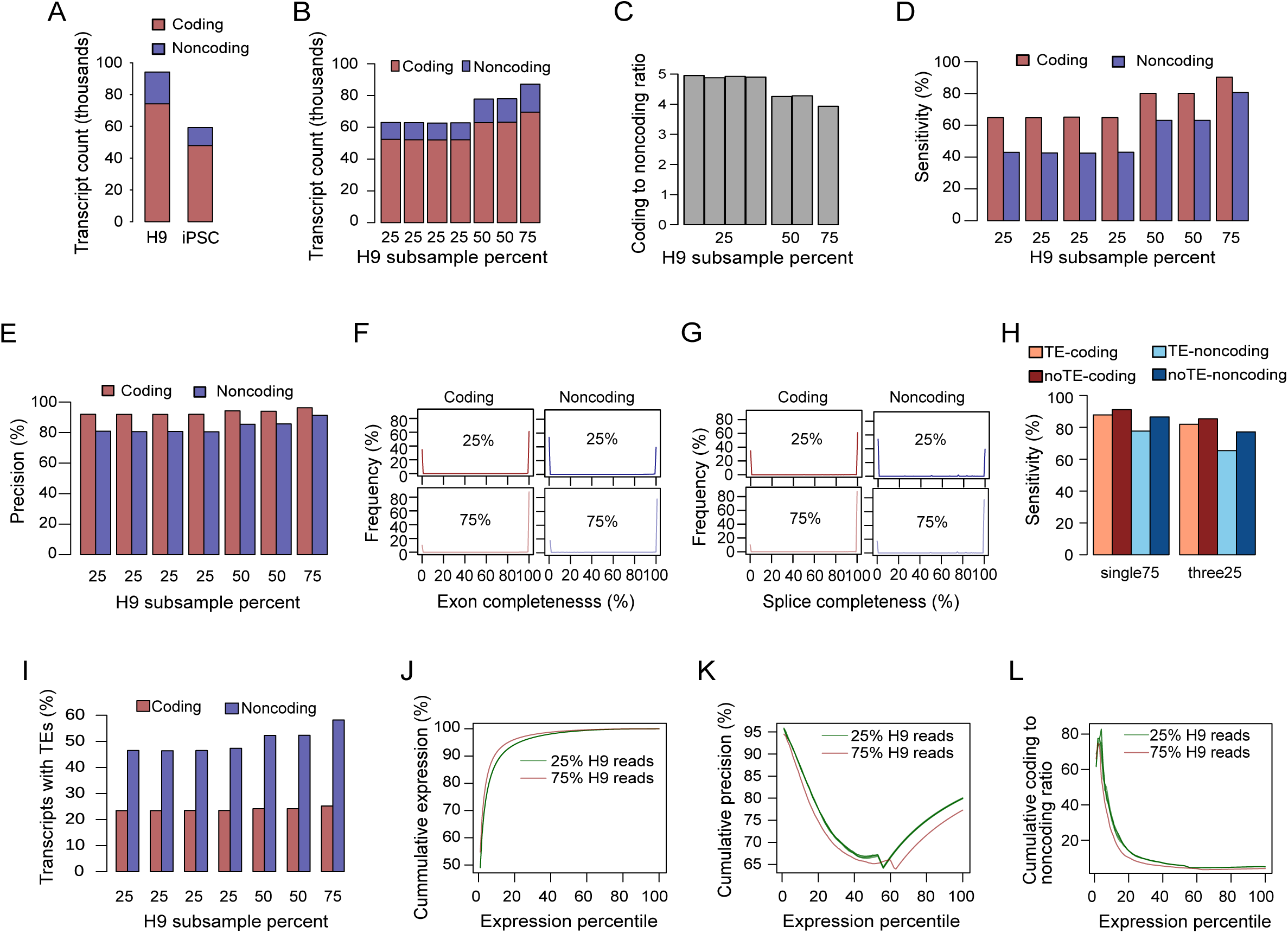
Effect of sequencing depth on the numbers and correctness of transcripts assembled from long-reads data. **A**. Transcript counts from iPSC and H9 human cell lines. 25% (n=4), 50% (n=2) and 75% of the H9 reads were independently sampled without replacements. **B.** Numbers of assembled transcripts from down-sampled H9 long-read data. Effects of down-sampling on coding to noncoding ratio **(panel C)**, sensitivity **(panel D)** and precision **(panel E)**. The transcript assembly from all the H9 long-read data was used as the reference for the computations of sensitivity, precision and completeness. The exon (**panel F)** and splice (**panel G**) completeness of the reference coding and noncoding transcripts in the 25% (n=4) and 75% (n=1) samples**. H**. The impacts of transposable elements inclusion on the sensitivity. Three replicates of 25% sampled long-reads were merged and compared to the assembly of 75% sampled long-read assembly. **I**. The proportions of TE-containing transcripts in the assemblies with various read depths. **J.** The cumulative expression of transcripts ranked by expression levels in H9 samples for 25% and 75% subsampled read assemblies. **K.** The cumulative precision of the expression-ranked transcripts for 25% and 75% subsampled read assemblies. **L.** The cumulative coding to noncoding ratio of the expression-ranked transcripts.

Because of the limited number of iPSC-sequenced long-read samples, we focused on only the deeper sequenced H9 cell line for the subsequent analyses. To assess the impact of long-read depth, H9 reads were randomly subsampled to 25% (four replicates), 50% (two replicates) and 75% (one replicate). Assemblies with a similar number of reads had similar numbers of transcripts (**Figure 4B**). We confirmed that the number of reads correlated with the number of assembled transcripts. It is important to note that the range of the number of reads in **Figure 3A** (10 – 200 million reads) is larger than in **Figure 4B** (∼6 – 19 million reads). We also established that the coding to noncoding ratio is higher in samples with fewer reads than in samples with more reads (**Figure 4C**). For sensitivity and precision, the transcript set assembled from the full H9 dataset reads was used as a reference. While the sensitivity was noticeably correlated with the number of reads (**Figure 5D**), the precision was not substantially affected (**Figure 5E**). This suggests that the increases in read depth led to the assembly of more reliable transcripts. Additionally, the comparison of the transcripts relative to the reference also shows two clear peaks in the exon (**Figure 4F)** and splice (**Figure 4G**) completeness. In both exon and splice completeness, the difference between coding and noncoding transcripts, and the effect of the number of reads was obvious. The results show that read depth affects the composition and integrity of the assembled transcripts.

**Figure 5.**
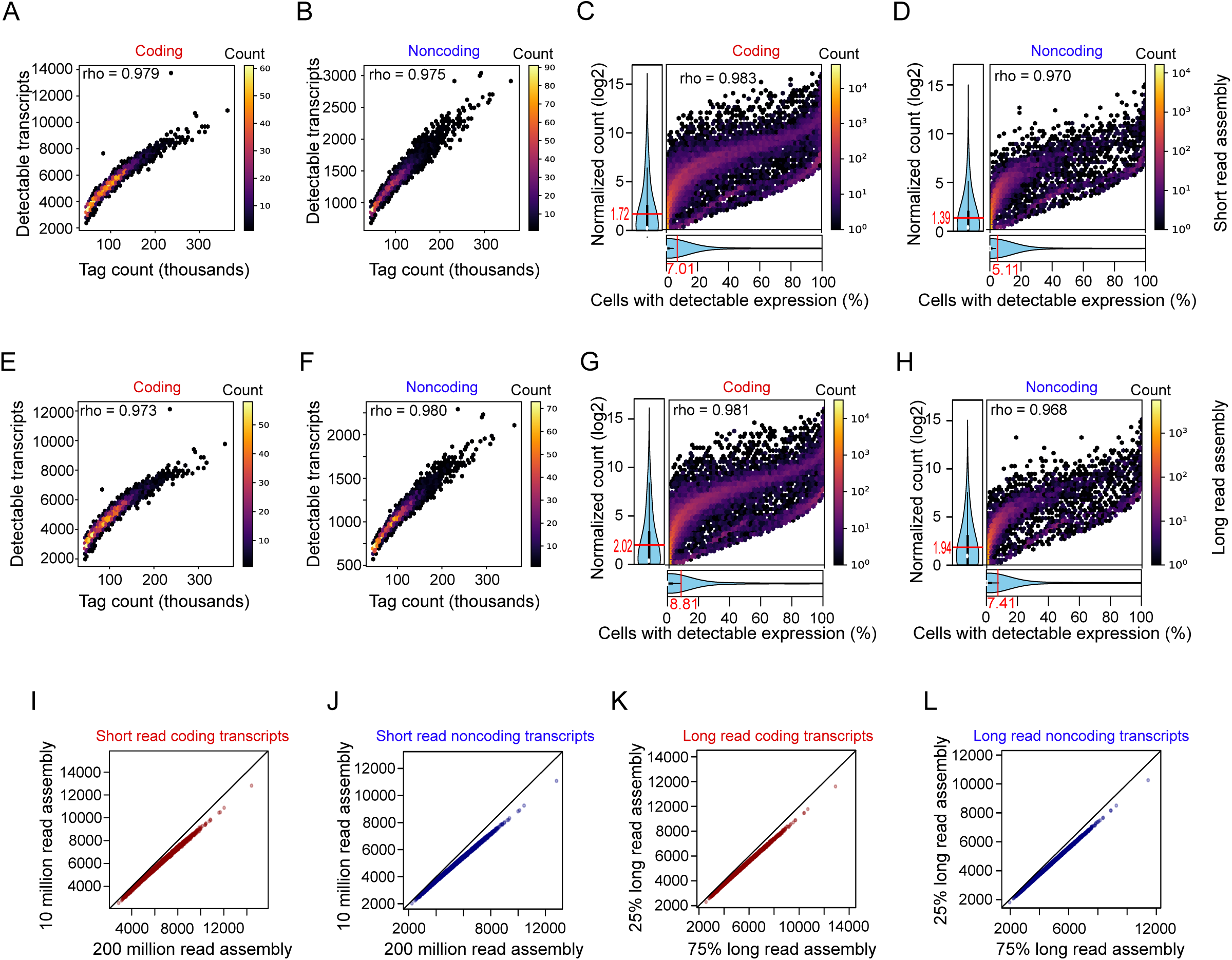
Single cell data reveal the heterogeneity of transcripts assembled from different data sources and read depths. The top 3,000 barcodes, corresponding to 3,000 cells were extracted from the alignment of c11/S0730 cell line. **Panels A-D** show the results from short-read transcript assembly using SRR597912 and SRR597895, and **panels E-H** show the results for the long-read H9 sample. The numbers of coding (**panel A**) and noncoding (**panel B**) transcripts assembled from short-read data are positively correlated to the tag counts of the barcodes. The normalized count of the coding (**panel C**) and noncoding (**panel D**) transcripts are positively correlated to the percentages of cells in which the transcript expressions are detected in sc-RNA-seq data. Positive correlations between the numbers of coding (**panel E**) and noncoding (**panel F**) transcripts assembled from long-reads and the sc-RNA-seq tag counts are shown. The normalized counts of the coding (**panel G**) and noncoding (**panel H**) transcripts from long-read assembly are positively correlated to the percentages of cells with detectable expression in sc-RNA-seq data. The means of the expression levels and percentages of cells with detectable expressions are shown in red fonts in panels **C**, **D**, **G** and **H**. The numbers of detectable transcripts in sc-RNA-seq data were compared for transcripts assembled from different read numbers of short-read (**panels I** and **J**) and long-read (**panels K** and **L**) data. For coding (**panel I**) and noncoding (**panel J**) transcripts, more transcripts assembled from 200 million reads were detected in sc-RNA-seq data than the transcripts assembled from 10 million reads. More coding (**panel K**) and noncoding (**panel L**) transcripts detectable from sc-RNA-seq data were found for the assembly from 75% long-read data than for the assembly from 25% long-read data. For **panels K-L**, each point represents a cell. The diagonals represent the points of equal numbers of transcripts from the compared assemblies.

We merged the three replicates of 25% subsamples (three25) and compared them with the 75% subsample (single75) in the context of the coding ability and presence of TE sequence fragments. The results showed that the assembly of coding transcripts generally tended to have higher sensitivity (**Figure 4H**). Also, TE presence tended to affect sensitivity. Interestingly, we found that single75 performed slightly better than three25, indicating that sequencing a larger number of replicates in long-reads might not be advantageous. It is important to note that the difference might be due to the sub-sampling procedure, as each replicate was sub-sampled with replacement. Because of the limited sampling space, there is a chance that some reads would be sampled in more than one replicate for the 25% sub-sample pseudo-replicates, whilst the 75% pseudo-replicate sub-sampling ensures that only unique reads were sampled. Interestingly, the proportion of TE-containing transcripts increased with depth in noncoding transcripts, but not in coding transcripts (**Figure 4I**), further highlighting the advantage of long-read data in detecting TE-containing transcripts, which are predominantly noncoding.

We also investigated the impact of transcript expression level on transcript assembly from long-read data. We ranked the transcripts by expression in the 25% and 75% samples. The analyses of cumulative expression show that the top 10% of the transcripts account for about 90% of RNAs in the 25% subsamples, and about 93% of the total RNAs in 75% subsamples (**Figure 4J**). The precision of transcript assembly also tended to vary across expression levels (**Figure 4K**). Mainly, the overall precision tended to be higher in the 25% samples, suggesting that fewer novel transcripts are retrieved from 25% samples. The coding to noncoding ratio varied greatly across the expression profiles (**Figure 4L**). The top 5% of the transcripts had a ratio of ∼80 while the ratio fell to <10 when the top 40% of the transcripts are considered. In most parts of the graph, 25% of samples had a higher ratio. These results reflect the effect of read depths and the composition of the transcripts greatly depends upon the expression thresholds set.

### Heterogeneity of the assembled transcripts at single cell levels

Unlike bulk short and long-read samples, single cell read coverage is typically too low for transcript assembly. As reads tend to be 3’ biased, the data is sparse and suffers from transcript dropout from individual cells, and analysis is generally gene-wise, rather than transcript-specific [53]. However, transcript assemblies from matching bulk samples can guide transcript analysis [54]. We reanalyzed the single cell RNA-seq (sc-RNA-seq) data from the c11 cell line [42]. The alignments were split into barcodes representing cells from which the RNAs were sequenced. First, we used the short-read assembly derived from the two largest short-read samples, SRR597912 and SRR597895 [50]. We checked the relationship between the barcode tag count and the number of transcripts with detectable expression in each cell. We found that the total sequence tag count was positively correlated with the number of coding (**Figure 5A**) and noncoding (**Figure 5B**) transcripts with detectable expression. Cells with more reads tended to have a larger number of detectable transcripts. As expected, the number of coding transcripts with detectable expression was more than the number of noncoding transcripts with detectable expression (**Figures 5A and B**). This suggests that the number of reads strongly affects single cell transcript quantification.

In short-read bulk samples, we established that coding transcripts tend to be expressed in more samples (**Figures 2G and 3H**) and at higher levels (**Figure S3D**) than noncoding transcripts. We therefore asked if there is a relationship between the expression levels and the percentage of cells with detectable expression. As expected, transcripts with higher expression levels tended to be expressed in more cells for coding (**Figure 5C**) and noncoding (**Figure 5D**) transcripts. Using the short-read transcript set, we also found that the expression level and the percentage of cells with detectable expression for the transcripts were higher in coding than in noncoding transcripts (**Figures 5C and D**). Short-read transcript assembly likely contains more transcriptional noise than long-read transcript assembly [42]. We therefore repeated the same analyses using the H9 long-read transcript assembly.

We confirmed the positive correlations between the barcode tag counts and the numbers of transcripts with detectable expression for both coding (**Figure 5E**) and noncoding (**Figure 5F**) transcript sets. We also confirmed positive correlations between expression levels and the percentage of cells with detectable expression for both coding (**Figure 5G**) and noncoding transcripts (**Figure 5H**). However, the expression levels of long-read assembled transcripts tended to be higher than the expression levels of short-read assembled transcripts (2.02 versus 1.72 for coding; 1.94 versus 1.39 for noncoding transcripts). Similarly, the percentages of cells with detectable expression when using the long-read transcript assembly (**Figure 5G and H**) were higher than the percentages for short-read transcript assembly (**Figures 5C and D**). These results suggest that sc-RNA-seq analysis guided by long-read assembly data is more reliable than short-read-guided analysis.

Finally, we investigated the effect of sequencing depth of the assembly on the number of transcripts with detectable expression in sc-RNA-seq data. For the short-read transcript assembly, we compared the transcript assemblies from 10 and 200 million reads. In both coding (**Figure 5I**) and noncoding (**Figure 5J**) transcript sets, the numbers of transcripts with detectable sc-RNA-seq expressions were higher in 200 million read assembly than in 10 million read assembly. We repeated the same analysis using shallow-depth (25%) and high-depth (75%) long-read data assemblies. The results show that the number of coding (**Figure 5K**) and noncoding (**Figure 5L**) transcripts from long-read assemblies with detectable sc-RNA-seq expressions were higher in high-depth assembly than in shallow-depth assembly. The results show that assembly from high-depth samples have more transcripts with detectable sc-RNA-seq expressions than the assembly from shallow-depth samples. These results further prove that shallow-depth samples might be missing more valid transcripts. Importantly, the results demonstrate that the effects of sequencing depth on transcript assembly can be detected in sc-RNA-seq data.

## Discussion

Previous studies have identified many differences between coding and noncoding transcripts [30,42,55–57]. Compared to coding transcripts, lncRNAs have been reported to be lowly expressed, lack open frames and known transcript features, and are evolutionarily less conserved across species, making ortholog discovery difficult [18]. In addition, noncoding RNAs tend to contain more TEs [42,58,59]. Consequently, they tend to be poorly annotated. Novel annotated transcripts tend to be dominated by noncoding transcripts [25,30,42,60]. Even for coding transcripts that have been relatively well-annotated, the precise number of genes remains controversial [24–26,61–63]. Because genes have different expression levels and sequence structure, the assembly of certain transcripts is more complicated than others [18,64,65].

In this study, we systematically investigated the effect of sequencing depth on transcript assembly for the human genome and particularly for hPSCs. Using short-read data from 671 tissues and cell types, we have established that the number of reads is positively correlated with the number of transcripts assembled. The correlation between the sequencing depth and the number of assembled transcripts was higher in noncoding transcripts, and confirmed in hPSC data, simulated short-read data, long-read data and sc-RNA-seq data. Additionally, we have established that the proportion of coding to noncoding transcripts decreases with increases in sequencing depth, indicating that noncoding transcripts benefit from deeper sequencing. The integrity of the assembled transcripts, measured by sensitivity, precision and transcript completeness, was also affected by the sequencing depth. All the data showed a positive correlation between sensitivity and sequencing depth, supporting the benefits of increased sequencing depth for improved transcript assembly. Although the precisions of the high-depth samples were not always higher than those of the shallow-depth sequenced samples, the loss in precision is not as rapid as the rise in sensitivity.

Several factors affect the extent to which the sequencing depth affects the number and accuracy of assembled transcripts. As stated above, noncoding transcripts tend to benefit more from higher sequencing depths. In fact, the assembly of lncRNAs in shallow-depth samples tended to be more stochastic, as they were correctly assembled in a smaller number of samples and expressed in fewer cells. Next, we found that transcripts in the GENCODE assembly tended to be correctly assembled, compared to novel hPSC transcripts. This discrepancy may be explained by limitations in GENCODE, particularly for noncoding transcripts as the GENCODE annotation is restricted in the assembly of certain transcript types [42, 48]. For examples, the latest GENCODE human release 39 (accessed on 1/21/2022) [27] had 53,009 human long noncoding transcripts corresponding to 18,811 genes. By comparison, NONCODEV6 database [66] had 173,112 human noncoding transcripts corresponding to 96,411 genes (accessed on 1/21/2022). Indeed, we found that GENCODE lncRNAs reach saturation quickly and that the assembly of TE-containing transcripts tended to be more difficult. As GENCODE is a pan-cell type annotation, it is likely missing cell type-specific lncRNAs. Indeed, there is growing evidence that noncoding RNAs, particularly those containing TEs, are highly divergent in different cell types [53, 54]. Another possible reason for the difficulty in assembling noncoding transcripts may be due to the lower apparent expression levels in bulk RNA-seq samples [29,41,42]. Highly expressed transcripts tend to be easily assembled, medium levels are more challenging to assemble, whilst low-expressed transcripts are assembled only if they have the support of reference transcripts. Finally, we assessed the impact of long-read sequencing, and comparing the results from the short-read and long-read assemblies, we demonstrated that long-read sequencing particularly benefits the assembly of TE-sequence containing noncoding transcripts, as previously suggested [41].

Our study shows that noncoding transcripts are difficult to assemble, compared to coding. This calls for caution as many noncoding transcriptional products have been described to be transcriptional noise [25,67,68]. Consequently, many studies have set expression threshold levels or used filters based on the number of expressing samples to remove unreliable transcripts [12,25,42,69–72]. However, threshold setting can substantially affect the nature and composition of the assembled transcripts. If the expression threshold is too high, more noncoding transcripts would be disproportionally filtered. Further, employing the same thresholds for both coding and noncoding transcripts might be inappropriate, as these classes of transcripts have different properties. For example, noncoding transcripts tend to have lower expression levels, be less stable, contain more TEs and are more likely to be localized in the nucleus [29,42,59,73,74]. Therefore, setting the same thresholds for coding and noncoding transcripts may bias against noncoding transcripts. While thresholds should be guided by downstream analyses, the nature of the transcripts should be considered while setting the thresholds. In conclusion, this study highlights the effects of the sequencing depth on transcript assembly, and how these impact transcript discovery and quantitation.

## Materials and Methods

### Data analyzed

The short-read data used in this study were from an in-house collection of publicly available data. The collection consisted of only paired-end short-read data with at least a 70% alignment rate to the human genome. The SRA accessions and the details of the data are presented in **Supplementary Table 1**. A total of 671 cells and tissue types were used for analysis. This collection included 150 human pluripotent cell samples that were previously analyzed in Babarinde et al. (2021). Two hPSC samples with the largest sequencing depths (SRR597912 and SRR597895 [50] were used for short-read sub-sampling. In addition to the short-read samples, four publicly available long-read samples (ENCFF688QGB, ENCFF272VSN, ENCFF954UFG and ENCFF251CBB) sequenced on the PacBio sequencing platform were also retrieved from the ENCODE project [75]. Single-cell data of c11/S0730 cell line (PRJNA631808) that were previously described [42] were also analyzed.

### Transcript assembly

The transcript assembly pipeline employed in this study is similar to the one used in Babarinde *et al.* For short-reads, the reads were first aligned to the hg38 version of the human genome using HISAT2 [76]. Transcripts were assembled from the short-read alignment using Stringtie [77, 78]. For long-read data, the fastq files were first converted to fasta format. Minimap2 [79] was then used for read mapping using the command below;

*minimap2 -t 30 -ax splice -uf --secondary=no -C5 -O6,24 -B4 Homo_sapiens.GRCh38.dna.primary_assembly.mmi $fasta > $out.sam 2> $out.err*

The alignments were then sorted with SAMtools [80] and Stringtie was used for assembly. After assembling, transcripts with no inferred strand or those shorter than 200 nucleotides were discarded [42].

### Coding potential classification

Transcript coding potential was measured using FEELnc [35]. Version 34 of human GENCODE protein-coding and long noncoding transcript sets were retrieved and used for training. FEELnc was run independently for each assembled set.

### Short-read data simulation

Two high-depth hPSC samples (SRR597895 and SRR597912) [50], which together had a total of 445 million paired-end reads, were used for short-read simulation. First, transcript assembly was done using the two samples. Transcripts with no strand information or those shorter than 200 nucleotides were discarded. The resulting transcripts were then used to build a reference for RSEM [49]. The counts, model and theta were estimated using *rsem-calculate-expression* with bowtie2 [81] alignment options. Finally, read simulations were done with *rsem-simulate-reads*. One replicate of 200 million reads, two replicates of 100 million reads, four replicates of 50 million reads, 10 replicates of 20 million reads and 20 replicates of 10 million reads were simulated.

### Long-read data sub-sampling

To investigate the relationship between the number of reads and assembly quality in long-reads, different fractions of long-reads from merged H9 alignments were sub-sampled using SAMtools *view -s* [80]. The investigated fractions were 0.25 (25%) with four replicates, 0.5 (50%) with two replicates and 0.75 (75%) with a single sub-sample. Independent transcript alignments were then carried out for each of the sub-sampled alignments. The unified transcript filtering and coding potential evaluation were then carried out for each assembly.

### Computation of transcript assembly integrity

We measured the integrity of transcript assembly with two parameters. Sensitivity measures the percentage of reference transcripts captured in the assembly. Precision measures the percentage of the assembled transcripts that are in the reference transcript set. Given the reference transcript set with *N* transcripts and a sample transcript assembly set with *n* transcripts, out of which *e* transcripts are correctly assembled as they are in the reference, sensitivity and precision are computed as follow.

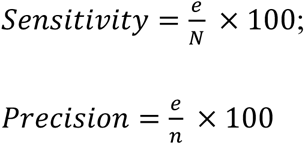

Further, transcript completeness was estimated for each transcript at the exon and splice levels. For a reference transcript, exon completeness was computed as the percentage of exons that are found in the closest assembled transcript. A transcript with 100% exon completeness has all the exons retrieved in the assembly. On the other hand, no exon of the transcript with 0% exon completeness was retrieved in the new assembly. Because exon overlap does not always mean splice overlap, we also computed splice completeness to highlight correct splice junctions in the assembly.

### Single-cell RNA-seq assembly

The FASTQ data from c11/S0730 cell line was processed and filtered as described in He et al., 2021. The alignments of the top 3,000 cells with the highest tag counts were retrieved and independently analyzed. The expression quantification was estimated from the barcode alignment using Stringtie [78]. The raw counts of the expression levels were computed using the Stringtie coverage per base and transcript length. For each sample, the expression matrix of the raw counts was then made for all the 3,000 barcodes. The raw counts were normalized with DESeq2 [82]. The expression levels of coding and noncoding transcripts for each assembly were independently estimated to compute the number of transcripts with detectable expression, the normalized tag counts and mean expression levels.

## Acknowledgements

This work was supported by the National Natural Science Foundation of China (31970589 and 32150710521), the Shenzhen Innovation Committee of Science and Technology (ZDSYS20200811144002008 to the Shenzhen Key Laboratory of Gene Regulation and Systems Biology), and the Stable Support Plan Program of the Shenzhen Natural Science Fund (20200925153035002). Supported by the Center for Computational Science and Engineering at the Southern University of Science and Technology.

## Author contributions

I.A.B. conceived the project, performed most of the analysis and wrote the manuscript. A.P.H. performed some of the analysis, revised the manuscript, funded and supervised the project.

## Tables and their legends

**Table S1:** Table summarizing the 671 short-read bulk paired-end RNA-seq samples. Columns include the sample description (the cell type or tissue as described in the study), the SRA or ERA accession number, and the total (unmapped read count).

## Supplementary figure legends

**Figure S1: Impacts of different counts of read depths on transcript assembly in multiple cells and tissues.** For each sample, the number of aligned reads and the number of reads assigned to transcripts were computed. **A**. Transcript count is positively correlated with the number of mapped reads**. B**. Transcript count is positively correlated with the number reads assigned to transcripts. **C**. The relationship between the number of reads and the number of mapped reads. **D**. The relationship between the number of reads and the number of reads assigned to transcripts. **E.** The relationship between the number of mapped reads and the number of reads assigned to transcripts. The difference between the observed and the expected numbers of coding (**panel F**) and noncoding (**panel G**) transcripts based on the loess prediction of all the 671 samples. Cells or tissues with at least 8 biological or technical replicates in the dataset are shown.

**Figure S2: Impacts of different counts of read depth on transcript assembly from short-read bulk RNA samples of human pluripotent stem cells. A.** For each sample, the number of aligned reads and the number of reads overlapping transcripts were computed. **A**. Transcript count is positively correlated with the number of mapped reads**. B**. Transcript count is positively correlated with the number of transcript-overlapping reads. The transcript-overlapping reads are more likely to have contributed to the transcript assembly**. C**. The relationship between the number of reads and the number of mapped reads. **D**. The relationship between the number of reads and the number of reads assigned to transcripts. **E.** The relationship between the number of mapped reads and the number of reads assigned to transcripts. **F**. The distribution of coding and noncoding transcripts in the unfiltered, GENCODE and filtered hPSC transcript assemblies.

**Figure S3: Impacts of read depth on transcript assembly in simulated human pluripotent stem cell reads. A.** Sensitivity of transcripts from merged reads. Different reads were merged such that the total number of reads was 200 million. For each category, aligned samples were merged before assembly. **B**. The precision of assembled transcripts from merged reads. **C**. The distribution of coding and noncoding transcripts in novel and GENCODE-annotated transcripts. Transcript assembly from merged SRR597912 and SRR597895 reads. The transcripts were then classified into “novel” or GENCODE-annotated based on the presence in GENCODE annotation. Novel transcripts can be a new isoform of a GENCODE-annotated gene or transcripts that do not overlap any known GENCODE gene. **D**. The expression levels of coding and noncoding transcripts in GENCODE and novel transcript sets. **E**. Transposable element presence in coding and noncoding transcripts from GENCODE-annotated and novel sets.

