## Supplementary figures and images for "The effects of sequencing depth on the assembly of coding and noncoding transcripts in the human genome"

### Supplementary Figures

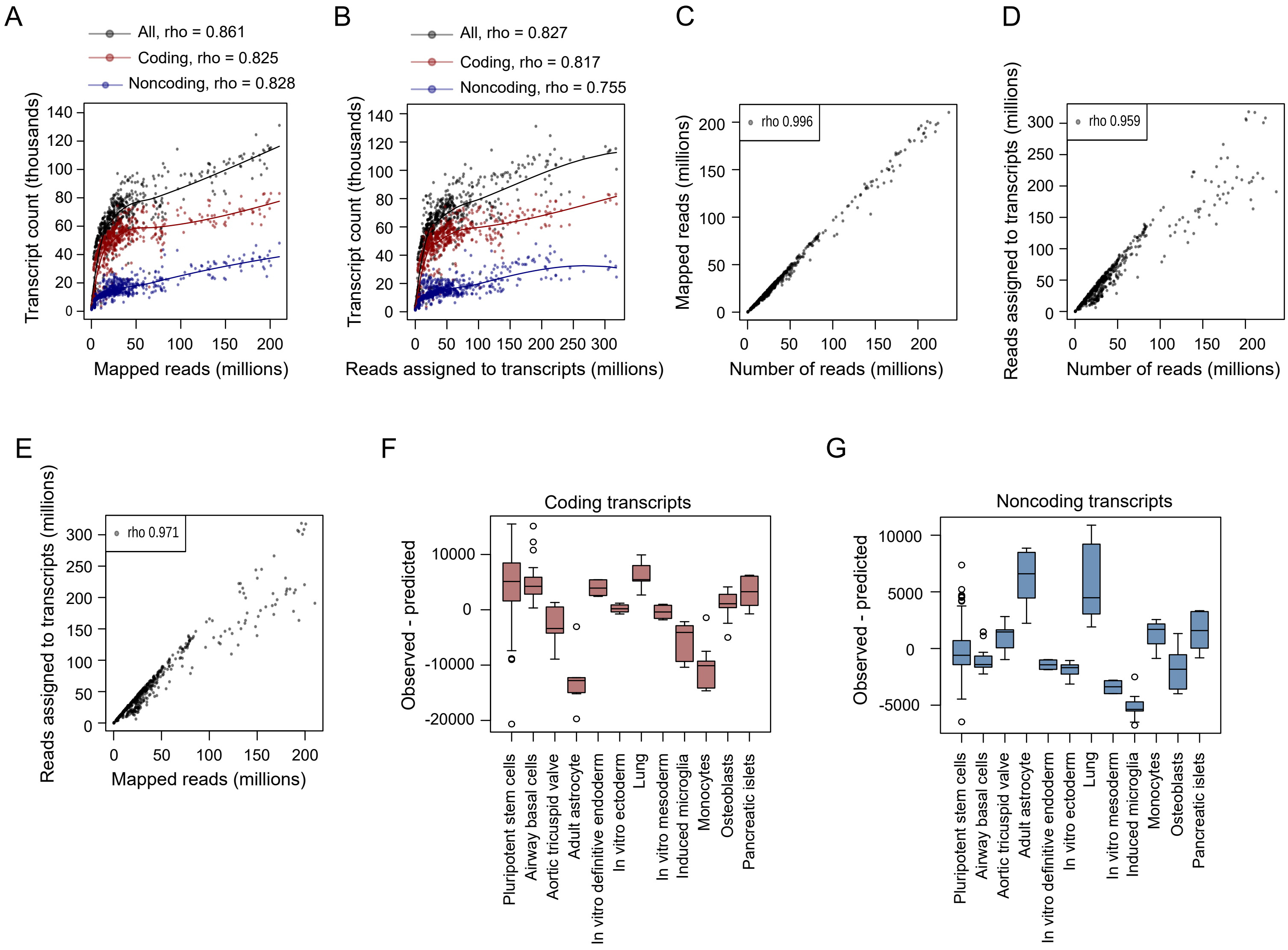

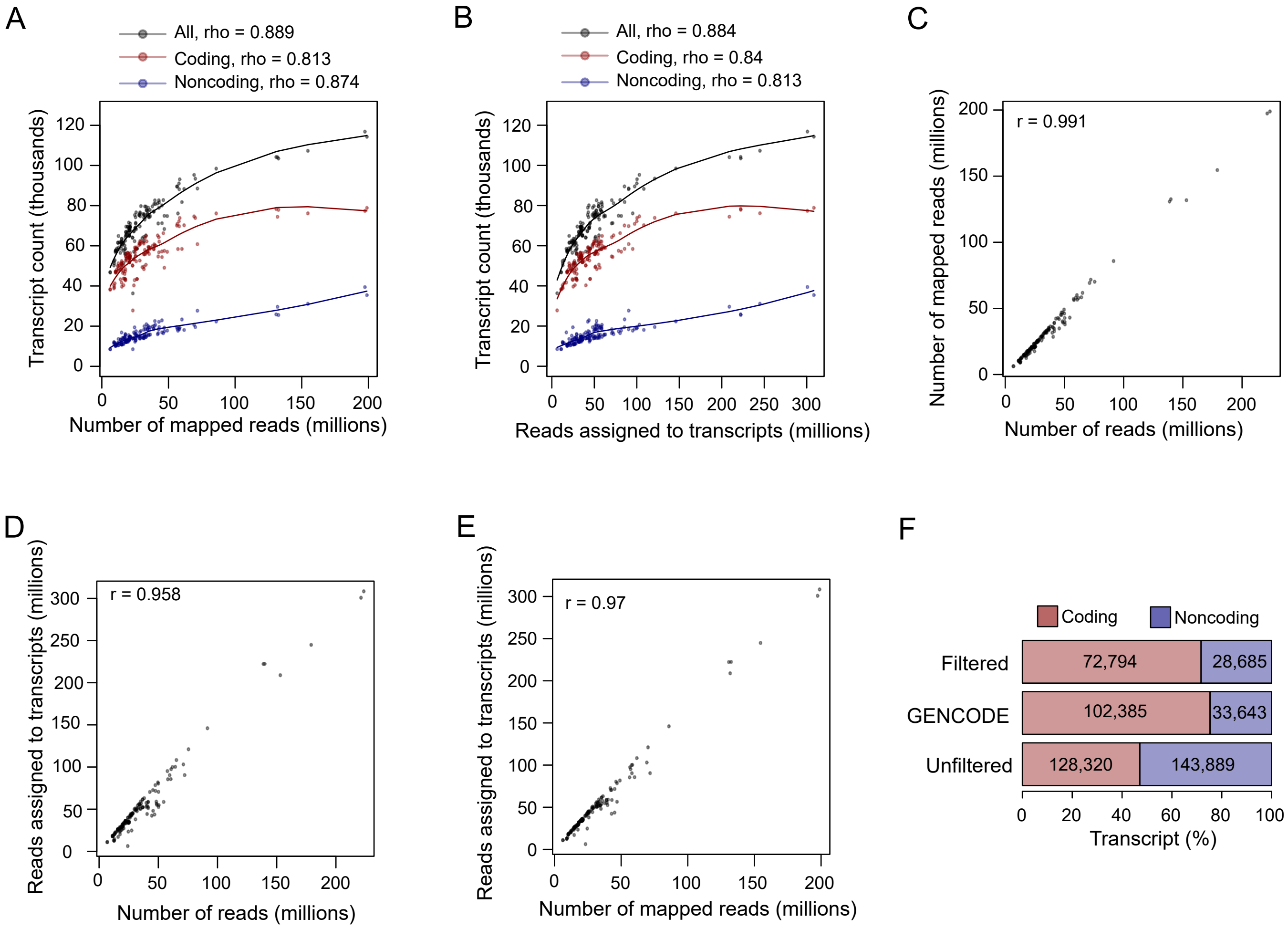

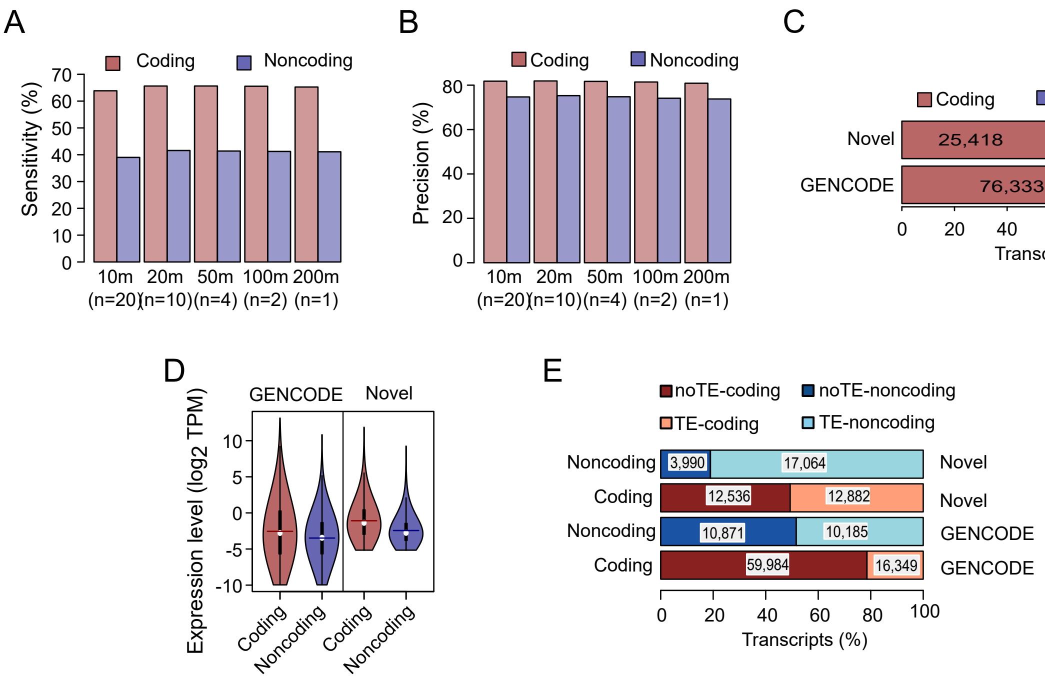
